# Mutational signatures are jointly shaped by DNA damage and repair

**DOI:** 10.1101/686295

**Authors:** Nadezda V Volkova, Bettina Meier, Víctor González-Huici, Simone Bertolini, Santiago Gonzalez, Harald Voeringer, Federico Abascal, Iñigo Martincorena, Peter J Campbell, Anton Gartner, Moritz Gerstung

## Abstract

Mutations arise when DNA lesions escape DNA repair. To delineate the contributions of DNA damage and DNA repair deficiency to mutagenesis we sequenced 2,717 genomes of wild-type and 53 DNA repair defective *C. elegans* strains propagated through several generations or exposed to 11 genotoxins at multiple doses. Combining genotoxin exposure and DNA repair deficiency alters mutation rates or leads to unexpected mutation spectra in nearly 40% of all experimental conditions involving 9/11 of genotoxins tested and 32/53 genotypes. For 8/11 genotoxins, signatures change in response to more than one DNA repair deficiency, indicating that multiple genes and pathways are involved in repairing DNA lesions induced by one genotoxin. For many genotoxins, the majority of observed single nucleotide variants results from error-prone translesion synthesis, rather than primary mutagenicity of altered nucleotides. Nucleotide excision repair mends the vast majority of genotoxic lesions, preventing up to 99% of mutations. Analogous mutagenic DNA damage-repair interactions can also be found in cancers, but, except for rare cases, effects are weak owing to the unknown histories of genotoxic exposures and DNA repair status. Overall, our data underscore that mutation spectra are joint products of DNA damage and DNA repair and imply that mutational signatures computationally derived from cancer genomes are more variable than currently anticipated.

## Introduction

A cell’s DNA is constantly altered by a multitude of genotoxic stresses including environmental toxins and radiation, DNA replication errors and endogenous metabolites, all rendering the maintenance of the genome a titanic challenge. Organisms thus evolved an armamentarium of DNA repair processes to detect and mend DNA damage, and to eliminate or permanently halt the progression of genetically compromised cells. Nevertheless, some DNA lesions escape detection and repair or are processed by error-prone pathways, leading to mutagenesis — the process that drives evolution but also inheritable disease and cancer.

The multifaceted nature of mutagenesis results in distinct mutational spectra, characterized by the specific distribution of single and multi-nucleotide variants (SNVs and MNVs), short insertions and deletions (indels), large structural variants (SVs), and copy number alterations. Studying comprehensive mutational patterns can yield insights into the nature of DNA damage and DNA repair processes. Indeed, large scale cancer genome and exome sequencing facilitated computational inference of more than 50 mutational signatures of base substitutions ^1,2^ in cancer cells. Some of these signatures, generally deduced by computational pattern recognition, have evident associations with exposure to known mutagens such as UV light, tobacco smoke, the food contaminants aristolochic acid and aflatoxins ^3–5^, or with DNA repair deficiency syndromes and compromised DNA replication. The latter include defects in homologous recombination (HR), mismatch repair (MMR), or DNA polymerase epsilon (POLE) proofreading. However, the etiology of many computationally extracted cancer signatures still has to be established, and it is not clear if these signatures have a one to one relationship to mutagen exposure or DNA repair defects.

The association between mutational spectra and their underlying mutagenic processes is complicated as mutations arise from various primary DNA lesions that include a multitude of base modifications and DNA adducts, single and double strand breaks, and intra- and inter-strand DNA crosslinks. This multitude of lesions is mended by numerous DNA repair pathways. Hence, there are at least two unknowns that contribute to a mutational spectrum: primary damage and DNA repair. Indeed, computational analyses of uterine cancers have identified distinct signatures associated with MMR and POLE exonuclease deficiency, respectively, and with the combination of both^6^. A non-additive signature change was also detected in *C. elegans* lines deficient for both the MMR factor *pms-2* and the polymerase epsilon subunit *pole-4*^*7*^. Due to its intrinsic error rate, polymerase epsilon incorporates mismatching bases during replication, most of which are repaired through the polymerase’s proofreading activity and by DNA mismatch repair. Hence, the signature of MMR deficiency is, in fact, a signature of polymerase errors that escaped its proofreading activity. When MMR and polymerase proofreading are lost simultaneously, the resulting mutational spectrum reflects intrinsic base incorporation errors and slippage events by polymerase epsilon. Analogous considerations apply when mutagenic lesions are inflicted by DNA damaging agents.

It is established that thousands of DNA lesions occur during a single cell cycle, even in healthy cells, and the number of lesions is further increased by exogenously applied genotoxic agents ^8^. However, only a tiny proportion of primary DNA lesions ultimately manifest as mutations, further highlighting the importance of the mechanisms that mend DNA damage in the genesis of mutational spectra. DNA repair capacity is also an important consideration for the apparent tissue specificity of some cancer-associated DNA repair syndromes; for example, inherited HR defects are commonly associated with breast and ovarian cancer, while Lynch syndrome characterized by MMR deficiency is primarily linked to colorectal cancer ^9^. DNA repair deficiencies play an important role in cancer development, permitting for more intense mutation acquisition. However, only a few of them have been assigned an associated mutational signature, namely biallelic MMR defects, specific variations of BER deficiencies, HR deficiency, and defective proofreading activity of the polymerase epsilon (POLE) ^1,2^.

Here we experimentally investigate the counteracting roles of genotoxic processes and the DNA repair machinery. Using *C. elegans* whole genome sequencing, we determine mutational spectra resulting from exposure to 11 genotoxic agents in wild-type and 53 DNA repair defective lines, encompassing most known DNA repair and DNA damage response pathways. Combining genotoxin exposure and DNA repair deficiency exhibits signs of altered mutagenesis, signified by either higher or lower mutation rates as well as altered mutation spectra. These interactions highlight how DNA lesions arising from the same genotoxin are processed by a number of DNA repair pathways, often specific for a particular type of DNA damage, therefore changing mutation spectra usually in subtle but sometimes also dramatic ways. In cancer genomes, except for rares cases such as the biallelic loss of NER, concurrent loss of MMR and POLE proofreading activity, or *MGMT* silencing combined with DNA alkylation by temozolomide, the loss of DNA repair capacity appears to only imprint moderate mutagenic effects. Despite these small effects, DNA repair defects are able to drive carcinogenesis, as evidenced by positive selection for missense and truncating mutations in DNA repair and damage response genes. In addition, considering that cancer transformation usually requires 2-10 independent mutations in driver genes and that the probability to independently mutate an increasing number of driver genes is a power of the mutation rate, even small increases in mutagenesis promote cancer development.

In summary, our results provide a comprehensive assessment of the phenomenon of DNA damage-repair signature interactions on a genome-wide scale. We confirm that such interactions are also imprinted in cancer genomes and provide an explanation why such cases are rare. The importance of minute mutagenic changes for cancer development underscores the need to understand the mutagenic changes impacted on affected genomes. At the same time, our data imply that mutational signatures derived from cancer genomes are more variable, and may not have a one-to-one relationship to distinct mutagenic processes.

## Results

### Experimental mutagenesis in *C. elegans*

To determine how the interplay between mutagenic processes and DNA repair status impacts mutational spectra, we used 54 *C. elegans* strains, including wild-type and 53 DNA repair and DNA damage sensing mutants (**Figure 1A**). These cover all major repair pathways including direct damage reversal (DR), base excision repair (BER), nucleotide excision repair (NER), mismatch repair (MMR), double strand break repair (DSBR), translesion synthesis (TLS), DNA crosslink repair (CLR), DNA damage sensing checkpoints (DS), non-essential components of the DNA replication machinery, helicases that act in various DNA repair pathways, telomere replication genes, apoptosis regulators and components of the spindle checkpoint involved in DNA damage response ^10–14^ (**Supplementary Table 1**). To inflict different types of DNA damage we used 12 genotoxic agents encompassing UV-B, X- and γ-radiation; alkylating agents such as the ethylating agent ethyl-methane-sulfonate (EMS), methylating agents dimethyl-sulfate (DMS) and methyl-methane-sulfonate (MMS); aristolochic acid and aflatoxin-B1, which form bulky DNA adducts; hydroxyurea as a replication fork stalling agent; and cisplatin, mechlorethamine (nitrogen mustard) and mitomycin C (which was only used in the wild-type) known to form DNA intra-and inter-strand crosslinks.

**Figure 1.**
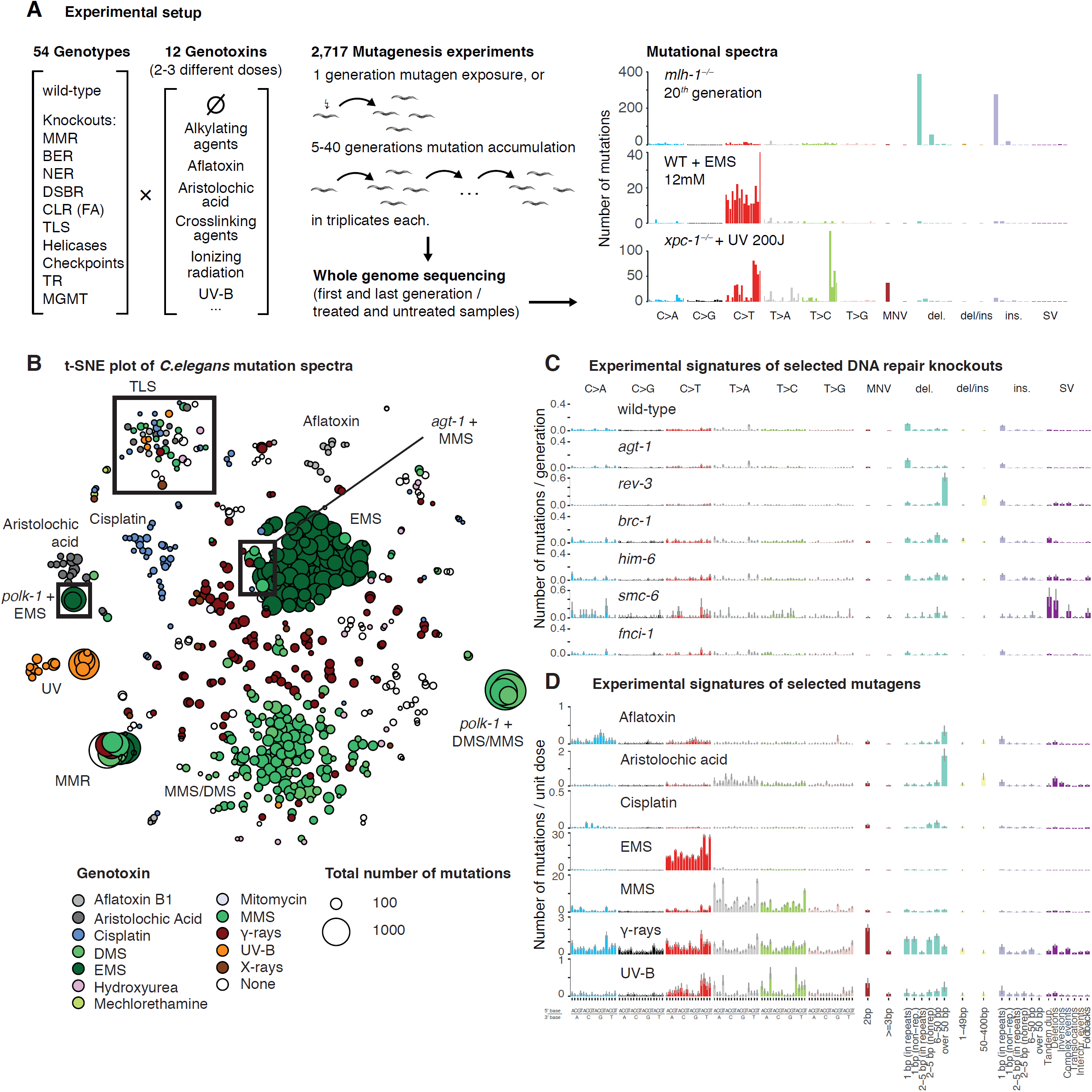
Experimental design of the study and data overview. A. Left: Wild-type and *C. elegans* mutants of selected genes from 10 different DNA repair pathways, including mismatch repair (MMR), base excision repair (BER), nucleotide excision repair (NER), double-strand break repair (DSBR), DNA crosslink repair (CLR), translesion synthesis (TLS), telomere replication (TR), and direct damage reversal by O-6-Methylguanine-DNA Methyltransferase (MGMT), were propagated over several generations or exposed to increasing doses of different classes of genotoxins. Genomic DNA was extracted from samples before and after accumulation or without and with treatment and sequenced to determine mutational spectra. Right: representative experimental mutational profile plots across 119 mutation classes: 96 single base substitutions classified by change and ‘5 and 3’ base context, di- and multi-nucleotide variants (MNV), 6 types of deletions of different length and context, 2 types of complex indels, 6 types of insertions and 7 classes of structural variants. B. 2D t-SNE representation of all *C. elegans* samples based on the cosine similarity between mutational profiles. Each dot corresponds to one sequenced sample, the size of the dot is proportional to the number of observed mutations. Colors depict the genotoxin exposures, no color represents mutation accumulation samples. Highlighted are selected genotoxins and DNA repair deficiencies. EMS, ethyl methanesulfonate; MMS, methyl methanesulfonate; DMS, dimethyl sulfate. C. Experimental mutational signatures of selected DNA repair deficiencies in the absence of genotoxic exposures. D. Experimental mutational signatures of selected genotoxin exposures in wild-type *C. elegans.*

To investigate the level of mutagenesis for each genotype in the absence of exogenous mutagens, we clonally propagated wild-type and DNA repair defective *C. elegans* lines over 5-40 generations in mutation accumulation experiments (**Methods, Figure 1A**) ^7,15^. In addition, to measure the effects of genotoxic agents, nematodes were exposed to mutagenize both male and female germ cells and mutational spectra were analysed by sequencing clonally derived progeny ^15^. Both approaches take advantage of the *C. elegans* 3 days generation time and its hermaphroditic, self-fertilizing mode of reproduction. Single zygotes provide a single cell bottleneck and subsequent self-fertilization yields clonally amplified damaged DNA for subsequent whole genome sequencing (**Figure 1A**). Experiments were typically performed in triplicates, with dose escalation for genotoxin treatments.

Overall, we analysed a total of 2,717 *C. elegans* genomes comprising 477 samples from mutation accumulation experiments, 234 samples from genotoxin-treated DNA repair proficient wild-type lines and 2,006 samples from interaction experiments combining genotoxin-treatment with DNA repair deficiency. The sample set harboured a total of 162,820 acquired mutations (average of ∼60 mutations per sample) comprising 135,348 SNVs, 937 MNVs, 24,308 indels and 2,227 SVs. Attribution of observed mutations to each genotoxin, to distinct DNA repair deficiencies, and to their combined action was done using a Negative Binomial regression analysis leveraging the explicit knowledge of genotoxin dose, genotype, and the number of generations across all samples to obtain absolute mutation rates and signatures, measured in units of mutations/generation/dose (**Methods**).

### Baseline mutagenic effects of DNA damage and repair deficiency in *C. elegans* and human cancers

#### Mutation accumulation under DNA repair deficiencies

Comparing mutations in the genomes of the first and final generations for the 477 mutation accumulation samples, we observed a generally low rate of mutagenesis in the range of 0.8 t0 2 mutations per generation similar to previous reports on 17 DNA repair deficient *C. elegans* strains ^15,16^ (**Supplementary Note, Supplementary Table 1**). Notable exceptions are lines carrying knockouts of genes affecting MMR, DSBR and TLS (**Figure 1C, Supplementary Note**). MMR deficient *mlh-1* knockouts manifested via high base substitution rates combined with a pattern of 1bp insertions and deletions at homopolymeric repeats, similar to previous reports in human cancers (cosine similarity *c*=0.85) ^7^. DSBR deficient strains carrying a deletion in *brc-1*, the *C. elegans* ortholog of the *BRCA1* tumor suppressor, exhibited a uniform base substitution spectrum and increased rate of small deletions and tandem duplications, features also observed in *BRCA1* defective cancer genomes (*c*=0.69 relative to SBS3) ^17^. TLS polymerase knockouts of *polh-1* and *rev-3* yielded a mutational spectrum dominated by 50-400bp deletions ^18^ (**Figure 1C, Supplementary Note**). Furthermore, *him-6* mutants, defective for the *C. elegans* orthologue of BLM (Bloom syndrome) helicase, and *smc-6* mutants, defective for the Smc5/6 cohesin-like complex, showed an increased incidence of SVs (**Figure 1C**). Experimentally derived mutational signatures for all 54 genetic backgrounds are summarized in **Supplementary Note** and **Supplementary Table 2**.

In cancers, deleterious mutations in genes associated with DNA repair or damage sensing are extremely common ^19^, yet they rarely are biallelic and incur complete deactivation (**Supplementary Materials, Supplementary Figure 1**). Similar to our *C. elegans* screen, DNA repair deficiencies alone do not seem to yield a strong change in mutation rates or spectra, with the exception of MMR, POLE exonuclease domain and HR defects (**Supplementary Figure 1**).

#### Mutagenesis under exposure to genotoxic agents

Visualizing the multi-dimensional mutation spectra for our entire *C. elegans* dataset in a 2D space based on their similarity to each other revealed some general trends: Genotoxins tend to have a stronger influence on the mutational spectrum compared to genetic background (**Figure 1B, Supplementary Figure 2A** for genotoxic effects in wild-type).

The methylating agents MMS and DMS, produced very similar mutation spectra with high numbers of T>A and T>C mutations (**Figure 1B, D**). Samples treated with the ethylating agent EMS were characterized by C>T mutations, somewhat similar to the mutational signature SBS11 observed in cancer genomes and attributed to the alkylating agent temozolomide (cosine similarity *c*=0.91)^2^, but different to the EMS spectrum in human iPS cells, possibly due to reduced metabolic activation (**Supplementary Figure 2B**)^20^. Bulky adducts created by aristolochic acid and aflatoxin-B1 exposure resulted in mutational spectra with typical C>A and T>A mutations, respectively, similar to those observed in exposed human cancers and cell lines (*c*=0.95 and *c*=0.92, respectively) ^4,21^. UV-B radiation induced characteristic C>T transitions in a YpCpH context (Y=C/T; H=A/C/T), similar to SBS7a/b (*c*=0.95 for C>T alone), but with an unexpected rate of T>A transversions in a WpTpA context (W = A/T; *c* = 0.69 for all variants), which might be caused by the higher frequency of UV-B-induced 6-4 photoproducts compared to daylight and possibly lower repair deficiency of these lesions in *C. elegans* ^22^. Ionizing radiation (X- and ɣ-rays) caused single and multi-nucleotide substitutions, but also deletions and structural variants, in agreement with the spectra of radiation-induced secondary malignancies ^23^ (*c* = 0.89 for substitution spectra). Lastly, cisplatin treatment induced C>A transversions in a CpCpC and CpCpG context, deletions, and structural variants in agreement with previous studies (**Figure 1 B,D**) ^15,20,24,25^. The overall spectrum of cisplatin exposure, however, seems to be strongly organism-specific (**Supplementary Materials**).

A summary of mutational spectra from *C. elegans* wild-type exposed to the 12 genotoxic agents is shown in **Supplementary Figure 2** and **Supplementary Table 2.** The general resemblance of signatures across organisms (**Supplementary Figure 2, Supplementary Materials**) reflects the high level of conservation of DNA repair pathways among eukaryotes. Observed discrepancies between species or different cell lines derived from the same species also provide insight, suggesting differences in genotoxin metabolism or DNA relative repair capacity.

### Interactions of DNA damage and DNA repair shape the mutational spectrum

The interplay between genotoxic agents and the repair machinery has not been systematically analysed *in vivo* by genome-wide analyses. Due to the experimentally controlled exposure time and doses, we were able to quantify how strongly a DNA repair deficiency alters mutational effects of a genotoxin on a genome-wide scale. We considered a combination of a genetic background and a mutagen as ‘interacting’ if the mutation rate of all or a class of mutations (eg. substitutions, indels or structural variants) changed relative to expectation that the observed mutations can be expressed as the sum of genotoxin spectrum in wild-type and the repair deficiency spectrum without genotoxin exposure (**Methods**).

Genotoxin-repair deficiency interactions were very common: In total, 88/196 (41%, at FDR < 5%) of combinations displayed an interaction between DNA repair status and genotoxin treatment involving 9/11 genotoxins and 32/53 genotypes (**Figure 2**, for a comprehensive list of effects see **Supplementary Note** and **Supplementary Table 3**). Usually interactions increased the numbers of mutations obtained for a given dose of mutagen up to 30x. Conversely, knockouts of TLS polymerases reduced the rate of point mutations for a range of genotoxins. While some interactions left the mutational spectrum largely unchanged, others had a profound impact on mutational spectra, indicating that a repair pathway is mending only a subset of lesions introduced by the same genotoxin (**Figure 2B)**.

**Figure 2.**
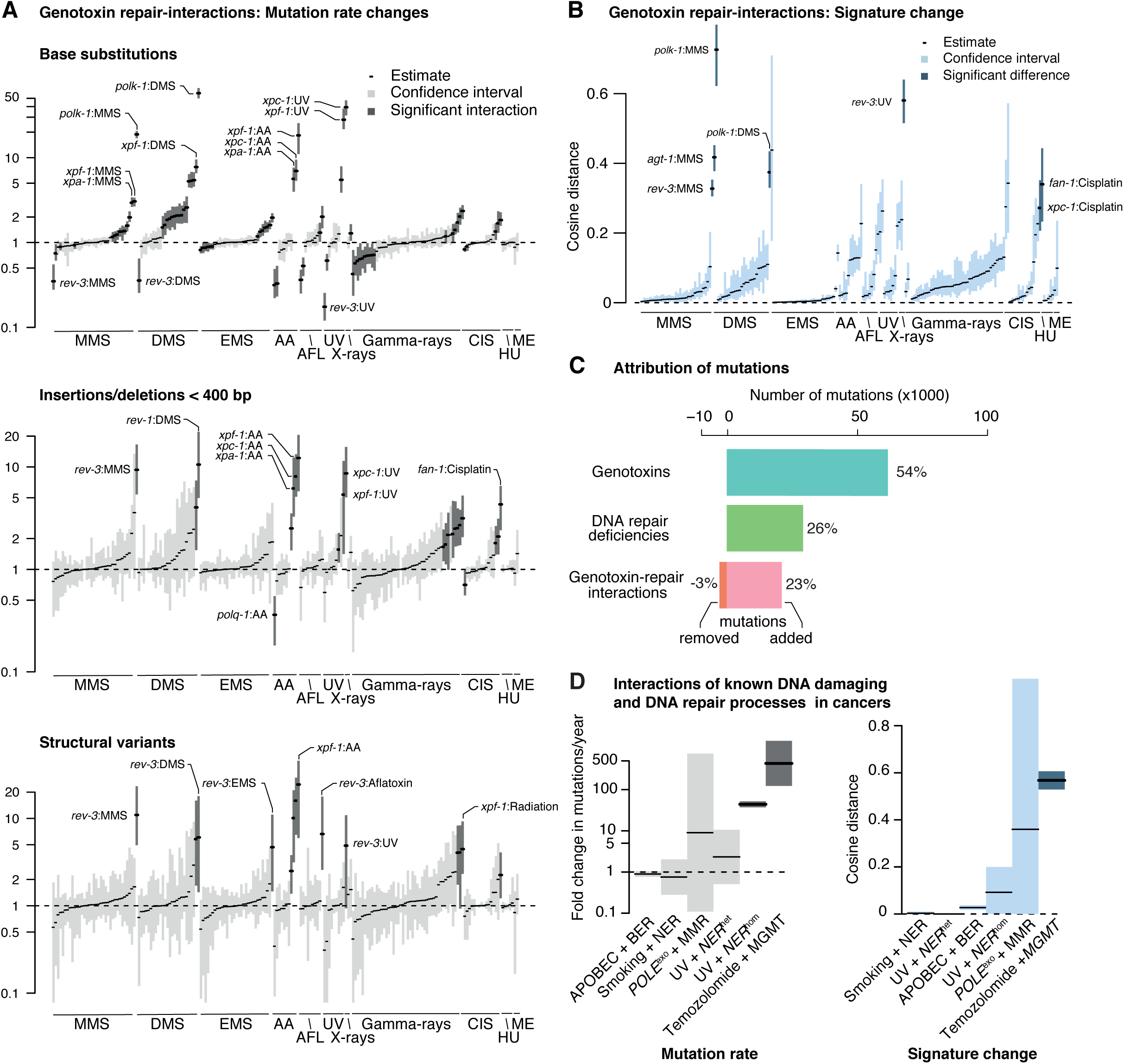
Widespread and diverse genotoxin-repair interactions in *C. elegans* and cancer. A. Estimated fold-changes between observed numbers of base substitutions relative to expected numbers based on additive genotoxin and DNA repair deficiency effects (upper panel). The same analysis conducted for insertions/deletions (middle panel) and structural variants (lower panel). A value of 1 indicates no change. Interactions with fold-changes significantly over or below 1 are shown in dark grey, the rest as light grey. Overall, about 37% of interactions produced a significant fold-change. AA, aristolochic acid. B. Changes in mutation signatures. Black lines denote point estimates. Interactions with cosine distances larger than 0.2 are shown with dark blue confidence intervals, others are shown in light blue. Around 10% of interactions produced a significant change in mutational spectra. C. Numbers of mutations attributed to different causes. Overall, about 23% of mutations in the dataset are attributed to genotoxin-repair interactions, including a reduction of –3% of mutations which would be expected under wild-type repair conditions. D. Summary of interaction effects of selected DNA repair pathway deficiencies with DNA damage-associated mutational signatures in human cancers. Left: Changes in age-adjusted mutation burden. Right: Change in mutational signatures.

To illustrate the overall magnitude of these interaction effects in our dataset, we estimate that of the 141,004 mutations we observed upon genotoxin exposure in DNA repair deficient strains, 26% were attributed to the endogenous mutagenicity of DNA repair deficiency genotypes independent of genotoxic exposure and 54% were attributed to genotoxic exposures independent of the genetic background. In addition, 23% of mutations arose due to positive interactions between mutagen exposure and repair deficiency leading to increased mutagenicity (**Figure 2C**). Finally, we found that 3% of mutations were prevented by negative interactions between genotoxins and DNA repair deficiency leading to reduced mutagenesis.

Applying a similar approach to a range of DNA repair deficient and proficient cancers with suspected genotoxic exposures, we were able to characterise several cases of repair-damage interactions leading to increased mutagenesis and/or altered signature, such as POLE^EXO^ and MMR, UV damage and NER, or APOBEC-induced mutagenesis and TLS (**Figure 2D, Supplementary Figure 3**). However, we found that such interactions were hard to detect due to unknown history of exposure and repair defects, and the detectable effects typically only revealed moderate effects.

#### Lesion-specific repair and mutagenicity of DNA alkylation

Many genotoxic agents, such as the alkylating agent MMS, inflict DNA damage on different nucleotides and residues thereof (**Figure 3A**). DNA methylation by MMS can lead to different base modifications, including non-mutagenic N7-meG, as well as highly mutagenic lesions such as N3-meA and O6-meG ^26^, which, taken together, produced a mutation rate of approximately 230 mutations/mM/generation in wild-type *C. elegans* (**Figure 3A**). However, the underlying mutagenic and repair mechanisms are very different: O6-meG is subject to direct reversal by alkyl-guanine alkyltransferases and tends to pair with thymine ^27^, whereas N3-meA can be mended by base or nucleotide excision repair ^28,29^. If unrepaired, N3-meA needs translesion synthesis polymerases to replicate across this base modification, which can both be error-free or result in the misincorporation of A or C opposite N3-meA ^30^. Hence, when different DNA repair components are unavailable, these lesions lead to different mutational outcomes.

**Figure 3.**
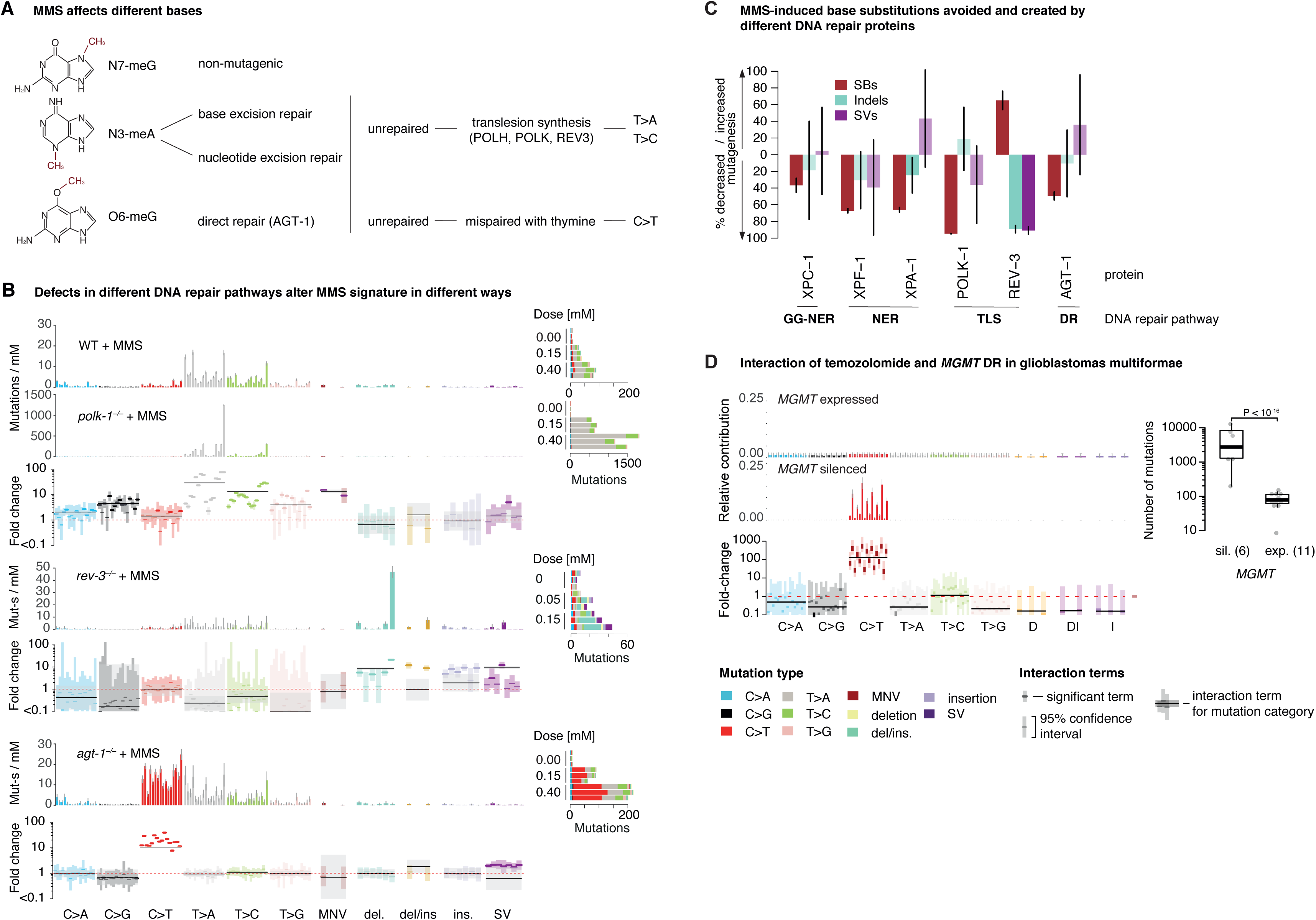
Knock-out specific signatures of MMS induced mutations reflect the activity of different DNA repair pathways. A. DNA repair and mutagenicity of different types of lesions induced by MMS exposure: The structures of non-mutagenic N7-methylguanine and mutagenic N3-methyladenine and O6-methylguanine are depicted. Methyl groups are indicated in red. B. Signatures of MMS exposure in wild-type, *polk-1, rev-3* and *agt-1* deficient *C. elegans*. Top: Mutation spectra of MMS exposure in wild-type and *polk-1*^−/–^ background. Error bars denote 95% confidence intervals. Below them is the plot of signature fold-changes per mutation type between the wild-type mutagen and interaction spectra. MMS treatment induced a 10-100x increase of T>A transversions in *polk-1* deficient background compared to wild-type. Darker lines denote the maximum a posteriori point estimates; faded color boxes depict 95% CIs. Point estimates and CIs are highlighted if different from 1. Middle: signature of MMS in *rev-3*^−/–^ background along with the respective fold-changes indicating the decrease in the number of substitutions, yet an elevated number of indels and SVs. Bottom: signature of MMS in *agt-1*^−/–^ background along with the respective fold-changes indicating a 10-fold increase in C>T mutations. Right: Dose responses for MMS in wild-type (upper part), *polk-1* deficient (middle part) and *agt-1* deficient (lower part) samples, measuring the total number of mutations (*x*-axis) versus dose (*y*-axis) for each replicate. Mutation types: MNV refers to multinucleotide variants, del/ins. to deletions with inserted sequences, SV to structural variants. TLS: translesion synthesis. DR: direct damage reversal. C. The fractions of mutations contributed or prevented by different DNA repair components compared to the mutations observed upon MMS exposure in wild-type *elegans*. Bars filled with more fainted color reflect non-significant changes. D. Mutational signature of temozolomide in glioblastomas with expressed (top) and epigenetically silenced *MGMT* (middle). Bottom: Mutation rate fold-change per mutation type. Right: Total mutation burden in temozolomide-treated samples with silenced and expressed *MGMT.*

Indeed, our analysis showed diverse changes in the MMS induced signature across DNA repair deficient backgrounds. In the absence of NER, mutation rate of adenine was elevated 1.5x in *xpc-1* mutants and 3x in *xpf-1* and *xpa-1* mutants without a change in the signature (**Figure 2A,B, Supplementary Figure 4A**), indicating that more adenine base modification underwent error-prone translesion synthesis. These fold-changes correspond to about 30% and 60% of mutations being prevented by XPC-1 and XPF-1/XPA-1 activity, respectively (**Figure 3C**). A similar trend for 1.5 or 3-fold increase in T>M mutations was observed for EMS exposure under NER deficiency (**Supplementary Figure 4B**). EMS mostly induces O6-meG lesions, with a small amount of N3-meA ^27^. In total, we estimate that NER prevents about 25-40% of mutations upon EMS exposure (**Supplementary Figure 4C**).

Deficiency of polymerase κ increased the total mutation rate upon MMS treatment 17x leading to approximately 3,800 mutations/mM/generation (**Figure 3B**). This increase also coincided with a distinct change in the mutational spectrum marked by the appearance of a prominent peak of T>A transversions in a TpTpT context (**Figure 3B**). In line with our expectations based on the role of TLS for tolerating N3-meA, polymerase κ deficiency yielded an approximately 10-100x higher rate of T>M mutations, especially in TpTpN and CpTpN contexts (**Figure 3B**). These figures indicate that Pol κ dependent TLS prevents 90-99% of DNA adducts caused by MMS treatment from becoming mutagenic (**Figure 3C**). A similar increase of T>M substitutions (at the rate of 10x) was observed in Pol κ deficient mutants upon treatment with EMS, corresponding to 50% of EMS-induced mutations being prevented by POLK-1 mediated error-free TLS (**Supplementary Figure 4B,C**). We postulate that in the absence of Pol κ, the bypass of alkylated adenines has to be achieved by other, error-prone TLS polymerases, leading to increased T>M mutagenesis, particularly in a TpTpT context.

One of the candidates for this error-prone TLS is *rev-3*, which encodes the catalytic subunit of TLS Pol ζ. In contrast to other combinations of exposure and DNA repair deficiency where the mutation rate was increased, the knockout of *rev-3/*Pol ζ partially suppressed MMS-induced base substitutions but increased the number of small deletions (**Figure 3B**), indicating that Pol ζ is an essential component for the bypass of alkylated bases. We estimated that about 60% of MMS mutations induced in the wild-type result from error prone repair synthesis conferred by Pol ζ (**Figure 3C**).

Combining MMS exposure with alkyl-transferase *agt-1* deficiency also led to a striking change in the MMS signature by increasing the C>T mutation rate from about 15 mutations/mM in the wild-type to over 200 in the mutant, while leaving the rate of T>M mutations unchanged (**Figure 3B**). This demonstrates that AGT-1 specifically reverses most O^6^-methylguanine adducts in an error free manner, thus acting as the functional *C. elegans* ortholog of the human O^6^-methylguanine DNA methyltransferase *MGMT*. Thus, the repair activity of AGT-1 prevents 50% of mutations which would otherwise be induced by MMS (**Figure 3C**). A similar, but weaker effect of *agt-1* deficiency was observed upon exposure to the ethylating agent EMS, leading to an increase in C>T transitions by a factor of 1.5, indicating that AGT-1 is also involved in, but less efficient at, removing ethyl groups, the latter result being consistent with reports measuring single locus reversion rates in *E. coli* ^*31*^ (**Supplementary Figure 4B,C**).

An even stronger interaction, which has already been therapeutically exploited, occurs between the human *agt-1* ortholog *MGMT* and temozolomide in temozolomide-treated glioblastomas (**Figure 3C**) ^32^. This is in good agreement with our experimental findings that the nature of mutation spectra detected upon EMS, DMS or MMS alkylation depends on the status of *agt-1* **(Figure 2A)**.

#### The majority of mutations induced by genotoxins are caused by error-prone translesion synthesis

Reduced mutagenesis in the absence of certain TLS polymerases is a wide-spread phenomenon ^33^. Error-prone TLS is a key mechanism used to tolerate several types of DNA damage, such as UV-induced cyclobutane pyrimidine dimers, which stall replication polymerases. Exposing wild-type to UV-B led to approximately 10 mutations/100J/m^2^, comprised mostly of C>T and T>C transitions (**Supplementary Figure 5A**). Paradoxically, the UV-B mutation rate was reduced 1.5-fold in *rev-3* mutants, mostly through a reduction in base substitutions, at the expense of a 4-fold increase in deletions longer than 50bp (**Supplementary Figure 5A**). To protect the genome from such mutations, which are likely to be deleterious, it is believed that TLS bypasses UV lesions at the cost of a higher base substitution rate ^33^. Quantifying the amount of mutations resulting from the activity of TLS polymerases REV-3/Pol ζ, POLH-1/Pol η and POLQ-1/Pol θ, we observed that 60-80% of the single base substitutions induced by aflatoxin, aristolochic acid, DMS, MMS and UV exposure could be assigned to the activity of REV-3/Pol ζ, which also prevents 40-80% of indels and structural variants (**Figure 4A**). A similar pattern was observed for POLH-1/Pol η, but, apart from aristolochic acid, to a lesser extent (**Figure 4A,B**).

**Figure 4.**
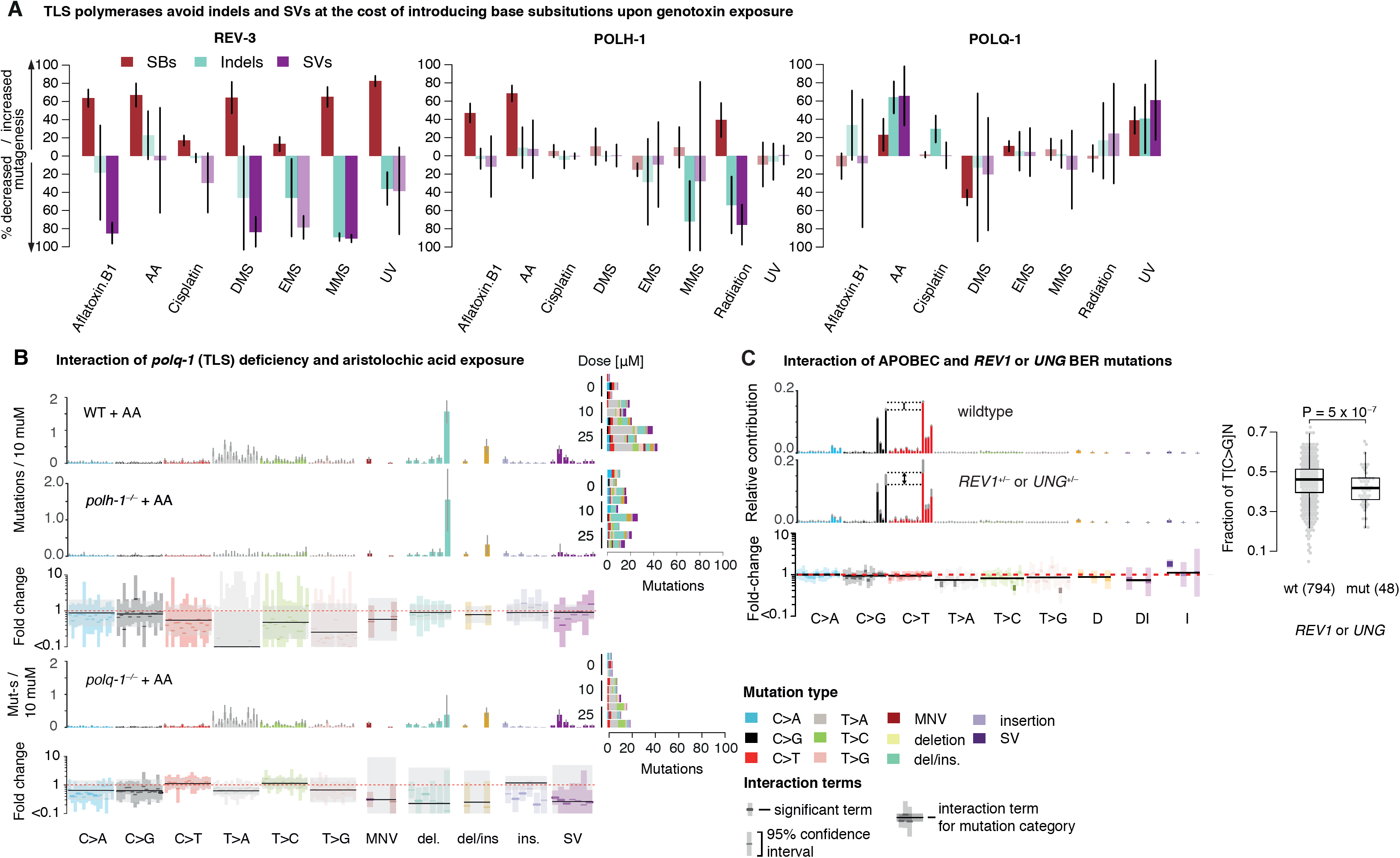
The majority of mutations observed upon genotoxin exposure result from error-prone TLS. A. The fractions of mutations caused or prevented by three TLS polymerases (REV-3, POLH-1 and POLQ-1) across genotoxin exposures. 60-80% of substitutions observed upon exposure are caused by REV-3 and POLH-1 in exchange for avoiding more deleterious indels and SVs. POLQ-1, on the contrary, introduces medium-sized deletions upon aristolochic acid and cisplatin exposure. B. Signatures of aristolochic acid exposure in wild-type and TLS polymerases η (*polh-1*) and θ (*polq-1*) deficient *C. elegans*. Same layout as in 3A. *polh-1* knockout yielded a 3-fold reduction in the number of base substitutions, while *polq-1* knockout led to nearly 5-fold reduction in the number of deletions compared to the values expected without interaction. C. Interactions of APOBEC-induced DNA damage with REV1 and UNG dependent BER. APOBEC mutation signatures in *REV1* and *UNG* wildtype (top) or deficient (middle) samples. Bottom: Fold-change of variant types. Right: Observed difference between C>G and C>T mutations in a TCN context.

An inverse phenomenon is observed upon treating *polq-1/*Pol θ deficient mutants with aristolochic acid (AA), where Pol θ causes more than half of indels and SVs (**Figure 4A,B**). Small deletions - between 50 and 400 bp in length - induced by AA in wild-type were absent in *polq-1* deficient mutants (**Figure 4B**). Breakpoint analysis of AA-induced deletions demonstrated an excess of single base matches, indicative of the *C. elegans* POLQ-1/Pol θ mediated end-joining ^18,34^ (**Supplementary Figure 5B,C, Supplementary Materials**). In the absence of POL-Q/Pol θ, replication-associated DSBs generated by persistent aristolactam adducts are likely to be repaired by HR, a slower but less error-prone pathway^35^.

A noteworthy example illustrating the roles of TLS synthesis in cancer is conferred by the APOBEC (apolipoprotein B mRNA editing enzyme, catalytic polypeptide-like) and error-prone TLS-driven BER. APOBEC deaminates single-stranded cytosine to uracil, which pairs with adenine during replication, leading to C>T mutations. Uracil is thought to be removed by Uracil DNA Glycosylase UNG; subsequent synthesis by error-prone TLS REV1 leads to C>G mutations ^36,37^. Indeed, a lack of C>G mutations has been observed in a cancer cell line with *UNG* silencing ^32,38^. We found that the evidence of *REV1* and *UNG* mutation or silencing contributing to APOBEC mutagenesis is weaker in human cancers, but nevertheless confirms the expected trend: On average, samples defective for *REV1* or *UNG* display an 8% decrease in C>G mutations (**Figure 4C**). However, *REV1* and *UNG* status at diagnosis only explains a small fraction of APOBEC mutations with low rates of C>G transversion, suggesting that other processes also contribute to these variants. Alternatively, the underlying signature change may not be detectable due to REV1/UNG mutation occurring late in cancer development compared to APOBEC overactivation.

#### Nucleotide excision repair mends the majority of genotoxic lesions

Nucleotide excision repair acts by excising a large variety of damaged bases, preventing up to 90% of mutations induced by different genotoxins, particularly those which induce damage to adenine or thymine (**Figure 5A**). After aristolochic acid exposure, *C. elegans xpf-1* mutants showed a 5-fold increase in mutation rate with a 20-fold increase in the number of 50-400 bp deletions, confirming that NER is crucial for the repair of bulky DNA adducts (**Figure 5B**). Aristolactam adducts occur on adenine ^39,40^, and the failure to exercise the modified based would lead to T>A changes or deletions. In contrast to previous reports on the lack of recognition of the aristolactam adducts by global genome NER (GG-NER)^41^, *xpc-1* mutants (which are defective for GG-NER) showed increased numbers of mutations upon aristolochic acid exposure, especially deletions in the range of 50-400 bps, similar to that in *xpf-1* mutants which are deficient in both GG-NER and transcription coupled NER (TC-NER) (**Supplementary Figure 6A**). Interestingly, only *xpf-1* (which is defective for both TC-NER and GG-NER) but not *xpc-1* deficiency led to an increase in the number of C>A mutations, suggesting that TC-NER maybe be more crucial to the repair of dG aristolactam adducts, which tend to be mispaired with adenine during Pol η mediated TLS ^42^.

**Figure 5.**
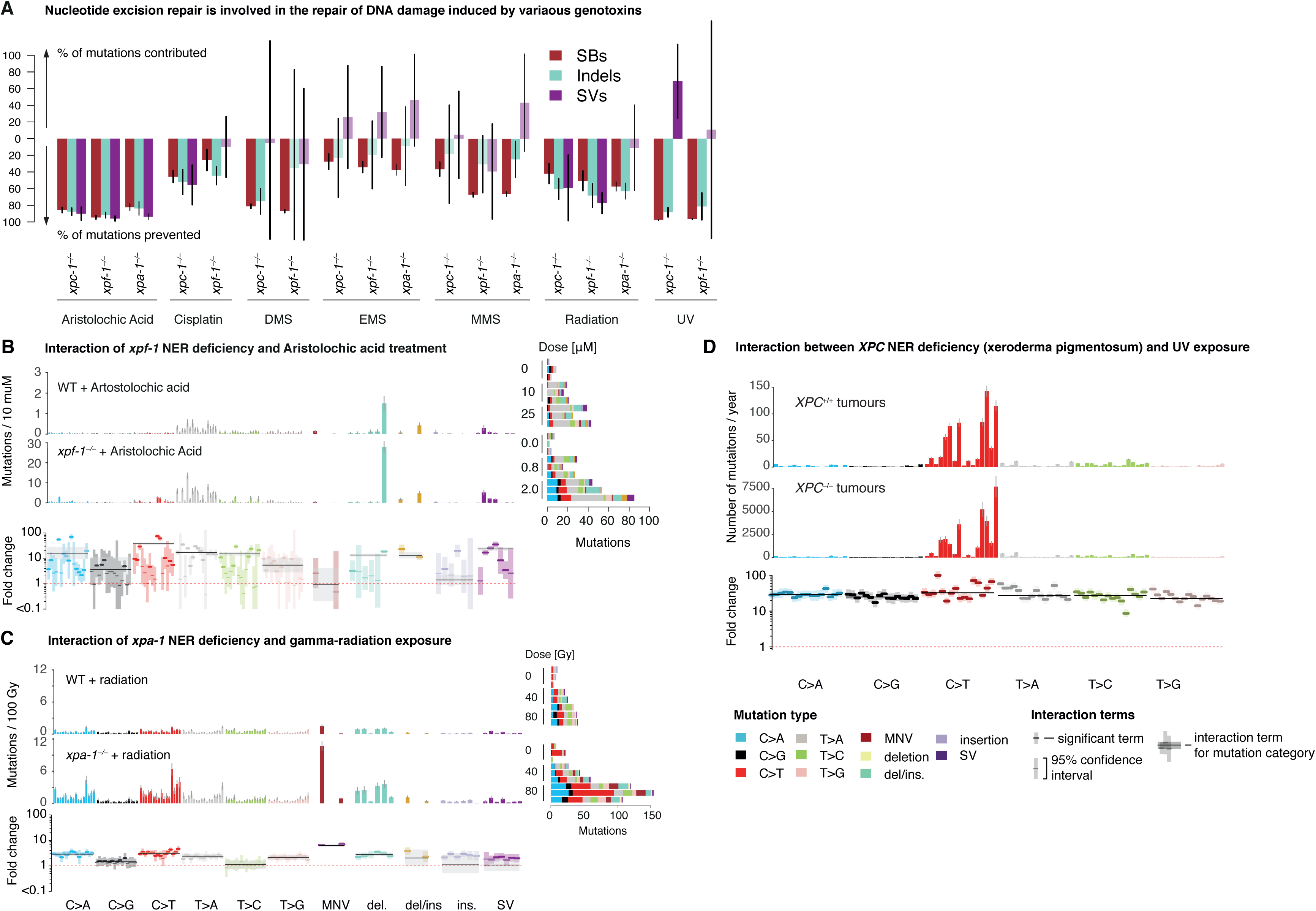
Nucleotide excision repair contributes to the repair of various genotoxins. A. Mutations contributed and prevented by different NER components for various genotoxins. NER prevents nearly 100% of mutations expected under AA and UV exposure, and plays a significant role in the repair of lesions introduced by every genotoxin tested. B. Signatures of aristolochic acid exposure in wild-type and *xpf-1* NER deficient *C. elegans*. The knockout of *xpf-1* leads to a 10-fold increase in the number of small deletions compared to the values expected without interaction. Note that scale bars are different in upper and middle panels. C. Signatures of gamma-irradiation exposure in wild-type and *xpa-1* NER deficient *C. elegans*. The knockout of *xpa-1* leads to a 2-fold increase in the number of mutations, and a 5-fold increase in TCN>TTN changes and dinucleotide substitutions in particular, compared to the values expected without interaction. D. Mutational signature of UV light in skin cSSCs samples from patients with functional (top panel) and mutated NER (XP patients, lower panel) expressed in mutations per year. The barplot underneath shows the fold-change per mutation type. The signature of UV exposure in the absence of GG-NER shifts towards more C>T mutations in ApCpT and TpCpW (W = A/T) contexts, and contributes about 30x more mutations than in GG-NER proficient tumours.

In contrast to the previous examples in which DNA repair deficiencies changed the mutational spectra induced by mutagens, knockouts of the NER genes *xpf-1* and *xpc-1* did not alter mutation spectra but uniformly increased the rate of UV-B induced mutagenesis by a factor of 20 and 32, respectively (**Figure 2A, Supplementary Figure 6B**). This uniform increase indicates that NER is involved in repairing the majority of mutagenic UV-B DNA damage including both single- and multi-nucleotide lesions. Interestingly, *xpf-1* and *xpa-1* knockouts also uniformly increased the mutation burden resulting from EMS and MMS alkylation by a factor of 2, indicating that NER also contributes to error-free repair of alkylation damage (**Supplementary Figure 4**). Further cases of genotoxin signatures being changed in NER mutants were observed following ionising radiation and cisplatin exposure (**Figure 5B, Supplementary Figure 6C**). In IR treated samples, *xpa-1* deficiency led to an up t0 10 fold increase in C>T changes as well as multinucleotide variants. Upon cisplatin exposure, NER defects increased the amount of C>A and T>A mutations, as well as dinucleotide substitutions possibly introduced during the bypass of cisplatin-induced crosslinks ^42,43^. In addition, in the *fan-1* mutant, defective for a conserved nuclease involved in ICL repair, cisplatin exposure led to a dramatic elevation of 50-500 base deletions without affecting base substitutions (**Supplementary Figure 6C**).

In humans, xeroderma pigmentosum (XP), a hereditary syndrome characterized by an extreme sensitivity to UV exposure and high risk of skin and brain cancers during childhood, is associated with biallelic inactivation of NER ^44^. Investigating the changes in the UV signature between 8 adult skin tumours and 5 tumours from XPC defective XP patients ^45^, we observed a relatively uniform 30-fold change in mutation rate per year in XP patients across base substitution types in line with findings in *C. elegans* (**Figure 5C**). At the same there is a mild shift in certain base contexts, with nearly 3 times more mutations acquired in NpCpT, but also TpCpD context. XPC deficiency inactivates GG-NER, which is possibly compensated by transcription coupled NER ^45^. It is thus possible that this shift in mutation spectrum reflects a sequence specificity of GG-NER repair efficiency.

Notably no effects were observed for NER variants in sporadic lung and skin cancers, although one might expect NER involvement in repairing bulky DNA adducts generated from tobacco smoke and UV light (**Supplementary Figure 3E-G**). Of note *ERCC2/XPD* NER mutations are also relevant in urothelial cancers, where they have been reported to produce a mild increase in the number of mutations attributed to COSMIC signature 5 ^46^.

## Discussion

Taken together, our experimental screen and data analysis show that mutagenesis is fundamentally driven by the counteraction of DNA damage and repair. A consequence of this interplay is that the resulting mutation rates and signatures vary in a number of ways. The systematic nature of the screen with multiple known doses of different genotoxins applied across a broad range of genetic backgrounds enabled us to precisely characterise how mutation patterns of genotoxic treatments change under concomitant DNA repair deficiency. Uniform mutation rate increases, without a change of mutational signature, suggest that a repair gene or pathway is involved in repairing all DNA lesions generated by the genotoxin. Examples of such interactions are NER deficiency combined with the exposure to UV-B damage, bulky DNA adducts, and alkylating agents. Conversely, mutation signatures change if divergent repair pathways are involved in repairing specific subsets of DNA lesions introduced by the same genotoxin. We illustrated this for alkylating agents, with the same adducts at different bases and residues being repaired by distinct pathways. In the case of DNA methylation, our data corroborate the notion that the mutagenicity of O6-methylguanine stems from mis-pairing with thymine, and is repaired by alkylguanine alkyltransferases, while N3-methyladenine stalls replication, which is resolved by translesion synthesis polymerases Pol κ and Rev-3/Pol ζ.

Our screen revealed that translesion synthesis plays an important and varied role in mutagenesis: Perhaps counterintuitively, error-prone TLS by Pol η (POLH-1) and ζ (REV-3) was found to *cause* the majority of base substitutions resulting from bulky adducts, alkylated bases, UV-B-induced damage and to a small extent cisplatin. Thus, knockouts of *rev-3* and *polh-1* resulted in reduced base substitution rates, but increased rates of large deletions and SVs, presumably due to replication stalling and fork collapse in the absence of TLS. Conversely, Polymerase κ (POLK-1) was found to prevent up to 99% of mutations induced by DNA alkylation, by performing largely error-free TLS across N3-meA and N3-etA. Lastly, Pol θ (POLQ-1) mediated deletions observed upon genotoxic exposures such as aristolochic acid provide a repair mechanism (TMEJ) for replication-associated DSBs ^18^. These findings imply that TLS – rather than the primary mutagenicity of genotoxic lesions – is an essential component of mutagenesis, and suggest a direction for further exploration in human tissue culture experiments. Based on the frequency of interactions involving various TLS polymerases, we suggest that a study of the genome-wide signatures of the errors introduced by those polymerases on different substrates may enhance the resolution of genotoxin signatures and provide the means to detect and target TLS polymerase deficiencies or overactivity in cancer genomes.

Our analysis of human cancers showed that, while non-silent mutations in DNA repair genes are common, bi-allelic inactivation of both copies generally required for loss of function appears to be a relatively rare phenomenon. Surprisingly, mutations in only a small number of DNA repair genes exhibited a measurable mutational phenotype on their own or in relation to other mutational processes. While the frequent lack of strong mutational signatures appears initially puzzling, there might be several explanations for this. If a DNA repair deficiency is acquired at a late stage, there may simply not have been enough time to accumulate substantial amounts of mutations. Moreover, the prevalence of small effects seems less surprising when seen through the lens of somatic evolution.

The evolutionary dynamics of cancer implies that small changes in mutation rates are likely to exert a large cancer-promoting effect: As cancer transformation requires between 2-10 driver gene point mutations ^47,48^, the chance to independently mutate multiple driver genes in the same cell becomes a power of the mutation rate ^49^. Indeed, such a relationship between the relative risk and mutation rates can be observed in a number of cancer predisposition syndromes, and smoking-related lung cancers also follow this trend (**Figure 6**). While MMR-deficient colorectal cancers have an ∼8-10 fold higher mutation burden than MMR-proficient carcinomas, inherited MMR deficiency increases colorectal cancer risk >115 fold ^49^. Similarly, while HR-deficient breast cancers only display a ∼3-fold higher mutation burden, this correlates with a 20-40 fold increased cancer risk for carriers of *BRCA1/2* mutations^50^. Even more so, XP patients display an 30-fold increased mutation rate ^45^, but have an approximately 10,000 fold increased rate of skin cancers ^44^. A further indication for the importance of losing the activity of repair genes during cancer development is the selective pressure for loss-of-function mutations over silent mutations (**Supplementary Materials, Supplementary Figure 3H**) ^47,51^. While the exact number of driver gene mutations remains unknown in each cancer type, an implication of these data is that small changes in mutation rates have a large impact on cancer risk, and conversely noticeable risk factors may derive from rather moderate mutagenic effects. These findings offer an explanation why genes with no overt detectable mutator phenotype may be positively selected in cancers. Furthermore, they underscore the need to accurately determine seemingly small and subtle mutagenic effects which are challenging to detect in cancer genomes, but are often found in our experimental screen (**Figure 2A,B**).

**Figure 6.**
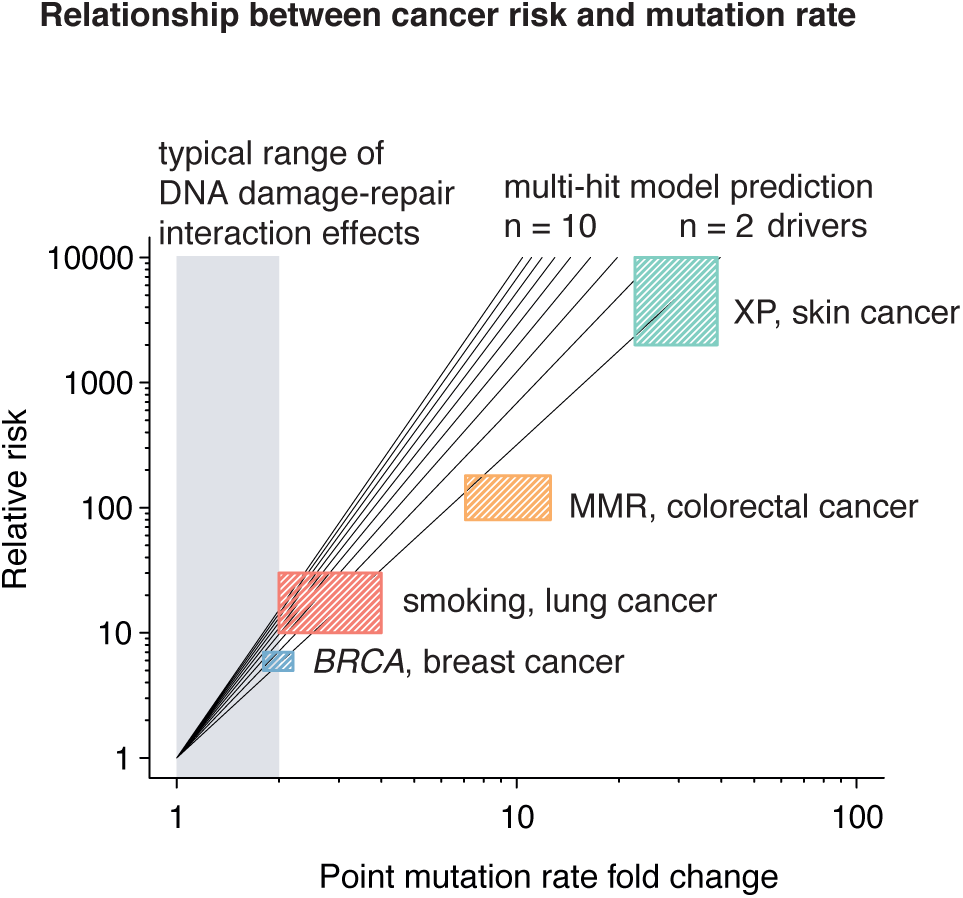
DNA repair deficiency drives cancer evolution. Relationship between relative cancer risk and point mutation rate change for different DNA repair pathway gene mutations. Black lines depict the expected power law depency in an evolutionary model with different number of driver gene mutations ^49^.

The analysis of mutational signatures derived from cancer genomes has gained much attention in recent years and dramatically deepened our understanding of the range and common types of mutation patterns observed in human cancer and normal tissues. These signatures revealed the mutagenic consequences of a number of endogenous and exogenous genotoxin exposures and signatures indicative of DNA repair deficiency conditions, which can be therapeutically exploited. Some mutational signatures derived from cancer genome analysis are very promising biomarker candidates, such as MMR deficiency for immunotherapy or HR deficiency for PARP inhibitor treatment. However, our study suggests that identifying signatures of other DNA repair deficiencies in cancer genomes may be more challenging as the resulting mutational signatures may be variable due to DNA-damage repair interactions, or simply manifest as an increased rate without a change in mutational signature as in the case of NER deficiency. Thus, to establish whether or not a DNA repair pathway is truly compromised, one might have to combine the analysis of mutational profiles with genetic and molecular testing of repair genes.

In order to learn from mutational signatures, it appears important to recognise the variable nature of mutagenesis due to the eternal struggle between DNA damage and repair. The high efficiency and redundancy of DNA repair processes removes a large fraction of genotoxic lesions and only damage left unrepaired results in mutations – either directly by mispairing or indirectly through error prone repair and tolerance mechanisms such as TLS. It should thus be kept in mind that the mutation patterns observed in normal and cancer genomes represent the joint product of DNA damage, repair, and tolerance.

## Acknowledgements

This work was supported by the Wellcome Trust COMSIG consortium grant RG70175, a Wellcome Trust Senior Research award (090944/Z/09/Z), a Worldwide Cancer Research grant (18-0644), and by the Korean Institute for Basic Science (IBS-R022-A1-2019) to AG. We thank the Mitani Lab funded by the National Bio-Resource Project of the MEXT, Japan, and the Caenorhabditis Genetics Center funded by the NIH Office of Research Infrastructure Programs (P40 OD010440) for providing strains. Nadezda Volkova is a member of Lucy Cavendish College, University of Cambridge. We thank Nuria Lopez-Bigas for critical comments on the manuscript.

## Data availability

Sequencing data are available under ENA Study Accession Numbers ERP000975 and ERP004086. Code for downstream analyses and figures can be downloaded from github.com/gerstung-lab/signature-interactions.

## Author contributions

NV, BM, AG and MG wrote the manuscript, which was approved by all authors. NV analysed all data with input from BM. BM performed mutation accumulation and mutagenesis experiments. VGH and SB performed mutagenesis experiments. SG and HV analysed somatic mutation data. FA and IM contributed to selection analysis. PC and AG conceived the study. AG and MG supervised the analysis.

## Methods

### Data generation

The experimental *C. elegans* data was generated as described previously ^15^, with strains and mutagen doses described in **Supplementary Table 1**. Hermaphrodites were treated with DNA damaging agents at the late L4 and early adult stages to target both male and female germ cells. Resulting zygotes provide a single cell bottleneck where mutations are fixed before being clonally amplified during *C. elegans* development and passed on to the next generation in a mendelian ratio. The baseline of mutagenesis in wild-type and DNA repair defective strains was determined by independently propagating 3-5 self-fertilizing hermaphrodites per genotype typically for 20 or 40 generations as described ^7^. For these mutation accumulation experiments, the P0 (parental) or F1 (1st filial) generation and the final generation were sequenced.

### Data acquisition

DNA from *C. elegans* samples were prepared as described previously ^15^. Samples were sequenced using Illumina HiSeq 2000 and X10 short read sequencing platforms at 100 bp paired-end with a mean coverage of 50x. TCGA human cancer data was taken from NCI GDC (https://gdc.cancer.gov, ^52^) and respective studies ^32,53–58^.

### Mutation calling

*C. elegans* samples were run through the Sanger Cancer IT pipeline including CaVEMan for SNV calling ^59^, Pindel for indel calling ^60^, and BRASS ^59^ and DELLY ^61^ calling consensus for structural variant identification. Mutation calls were filtered as described ^15^. The clustering and classification of SVs was constructed based on ^62^ as described in **Supplementary Methods**. Additionally, duplicated mutations across samples were removed (**Supplementary Methods**). Mutation counts in all samples are listed in **Supplementary Table 1**.

Variants were classified into 96 SNV classes (6 base substitution types per 16 trinucleotide contexts), 2 MNV classes (dinucleotide variants and longer variants), 14 indels classes (6 classes of deletions based on deletion length and local context (repetitive region or not), similar for insertions, and two classes for short and long complex indels), and 7 SV classes (complex variants, deletions, foldbacks, interchromosomal events, inversions, tandem duplications and intrachromosomal translocations).

Data visualisation was performed using *t*-SNE ^63^ based on the cosine distance of the 119-dimensional mutation spectra, averaged across replicates.

### Extraction of interaction effects in experimental data

For each sample *i* = 1, …, 2721 and mutation class *j* = 1, …, 119 we counted the respective number of mutations per sample *Y* ^*ij*^. Using matrix notation, the counts *Y* ∈ ℕ^2721 × 119^ were modeled by a negative binomial distribution,

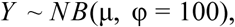

with scalar overdispersion parameter ϕ = 100 selected based on the estimates of overdispersion in the dataset, and matrix-variate expectation 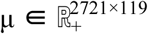. The basic structure of the expectation is

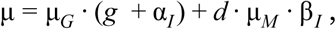

where 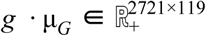 describes the expected mutations due to the genotype *G* in the absence of a genotoxin after *g* generations, while 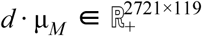 denotes the expectation for mutagen *M* at dose *d* in wild-type conditions. The terms 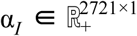 and 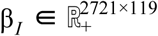 model how these terms interact (more details follow below). Here the dot product “·” is element-wise and we use the convention that a dimension of length 1 is extended by replication to match the length of the corresponding dimension of other factor in the product.

For α_*I*_ = 0, expected number of mutations for genotype G in wildtype has the structure

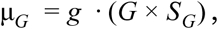

where 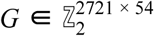 is the binary indicator matrix for denoting which one of the 54 genotypes a given sample has (*G*^*ij*^ = 1 if sample *i* has genotype *j*, otherwise *G*^*ij*^ = 0). The matrix 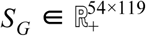 denotes the mutation spectrum (i.e. signature) across the 119 mutation classes for each of the 54 genotypes in the absence of genotoxic exposures. To provide a more stable estimate for genotype signature, the prior distribution for *S*_*G*_ is derived from 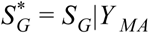, where *Y* _*MA*_ is the matrix of counts coming from mutation accumulation samples only, and 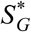 has a log-normal prior distribution 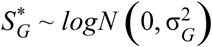 iid with a scalar variance 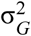. The vector 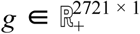 denotes the generation number in every sample, *g* = *g*(*N*), adjusted for the chances of fixation after *N* generations given the 25%-50%-25% probability of each mutation becoming homozygous, remaining heterozygous, or being lost in each generation. Similarly, the expected mutations for a given mutagen at dose 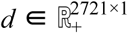 reads

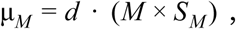

with 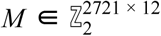 being the indicator for the 12 mutagens used in this screen and 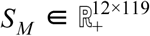 being their signatures in wildtype, with per-column prior 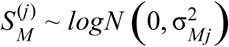, where the variances 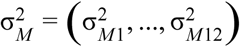 have prior distributions 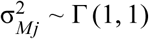 iid. Thus, the matrix-variate expected number of mutations can be written as

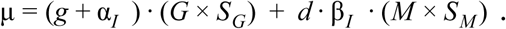

The vector-variate interaction term α_*I*_ measures how the mutations expected for genotype *G* may uniformly increase under mutagen exposure and is taken to be linear with respect to the mutagen dose,

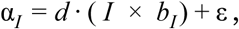

The symbol 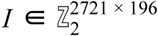 is the binary indicator matrix for each one of the 196 interactions tested in the screen multiplied by the interaction rate 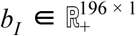, which measures how much the mutation signature of the genetic background increases upon genotoxic exposure with dose *d*. The interaction term is modelled to have a prior exponential distribution *b*_*I*_ ∼ *Exp* (1) iid.

Lastly ε is a random offset to allow for a possible divergence of the genotypes between experiments, adding εμ_*G*_ mutations with the same spectrum as in the absence of a mutagen in a dosage-independent fashion. The random offset is modelled as ε = (*J* × *a*_*J*_) where 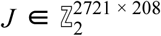 is an extended binary indicator matrix for each one the 196 interactions and 12 wild-type exposure experiments, with value 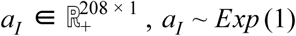 iid.

In addition to this scalar interaction, the wild-type spectrum of the mutagen *S*_*M*_ may change by the factors 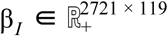 measuring the fold-change of each mutation type which is expressed as

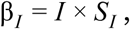

where 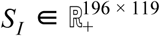 is the fold-change for each mutation class in a given interaction.

The prior distributions for each of the 119 subclasses of mutations in *S*_*I*_ are calculated in groups of 11 main mutation categories (6 types of SNVs, MNVs, 3 types of indels, SVs), since the numbers for individual mutation types were sometimes very small. The prior was defined using 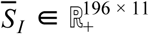 from an analogous model as described above, but applied only to observed mutation counts summed into 11 main mutation categories 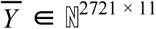. We used a Laplace prior 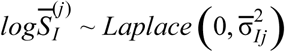, first to calculate 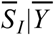. The posterior expected value 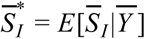 was chosen as the prior expectation for the 119-dimensional mutation effects 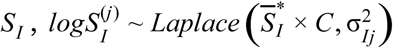 where *C* is the matrix spreading mutation category value across the corresponding individual mutation classes, and the variances 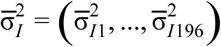 and 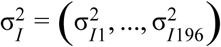 were assumed to come from Γ (1, 1) iid.

Bayesian estimates of the parameters *S*_*G*_, *S*_*M*_, *S*_*I*_, *a*_*J*_, *b*_*I*_ and hyperparameters 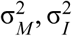 were calculated via Hamiltonian Monte Carlo using the ‘greta’ R package (http://CRAN.R-project.org/package=greta) (**Supplementary Methods**). Mean estimates along with their credible intervals may be found in **Supplementary Tables 2-3**.

Subsequent estimates of the fraction of mutations contributed or prevented by different DNA repair components upon genotoxin exposures were acquired as 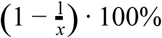 for positive interactions and as (1 − *x*) · 100% for negative interactions, where *x* is the fold-change in the mutation rate induced by the respective interaction compared to the exposure in wild-type.

### TCGA/PCAWG human cancer data analysis

The susceptibility of a DNA pathway to alteration was defined as having altered expression or high impact mutations in relevant genes (**Supplementary Methods, Supplementary Table 4**). The interaction effect was then estimated using Hamiltonian Monte Carlo sampling ^64^ for the following model for matrix of mutation counts in *N* samples with *R* mutation classes *Y* ∈ *Mat*_*NxR*_(ℕ ⋃ {0}):

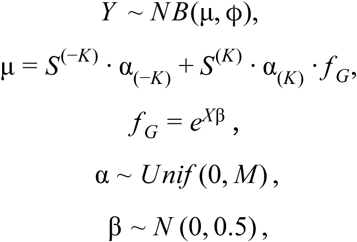

where α is the matrix of exposures (number of mutations assigned to each signature), *S* is a matrix of *K* signatures (each column is a signature, and is normalized to sum to 1), and *f*_*G*_ is the interaction factor which alters the signature. It consists of *X* - a binary matrix of labels, and β which is a matrix of spectra of the interaction effects. *M* is the maximal number of mutations per sample in the dataset. The overdispersion parameter ϕ = 50 was chosen based on variability estimates across all cancer samples.

## Supplementary Figures

**Supplementary Figure 1.**
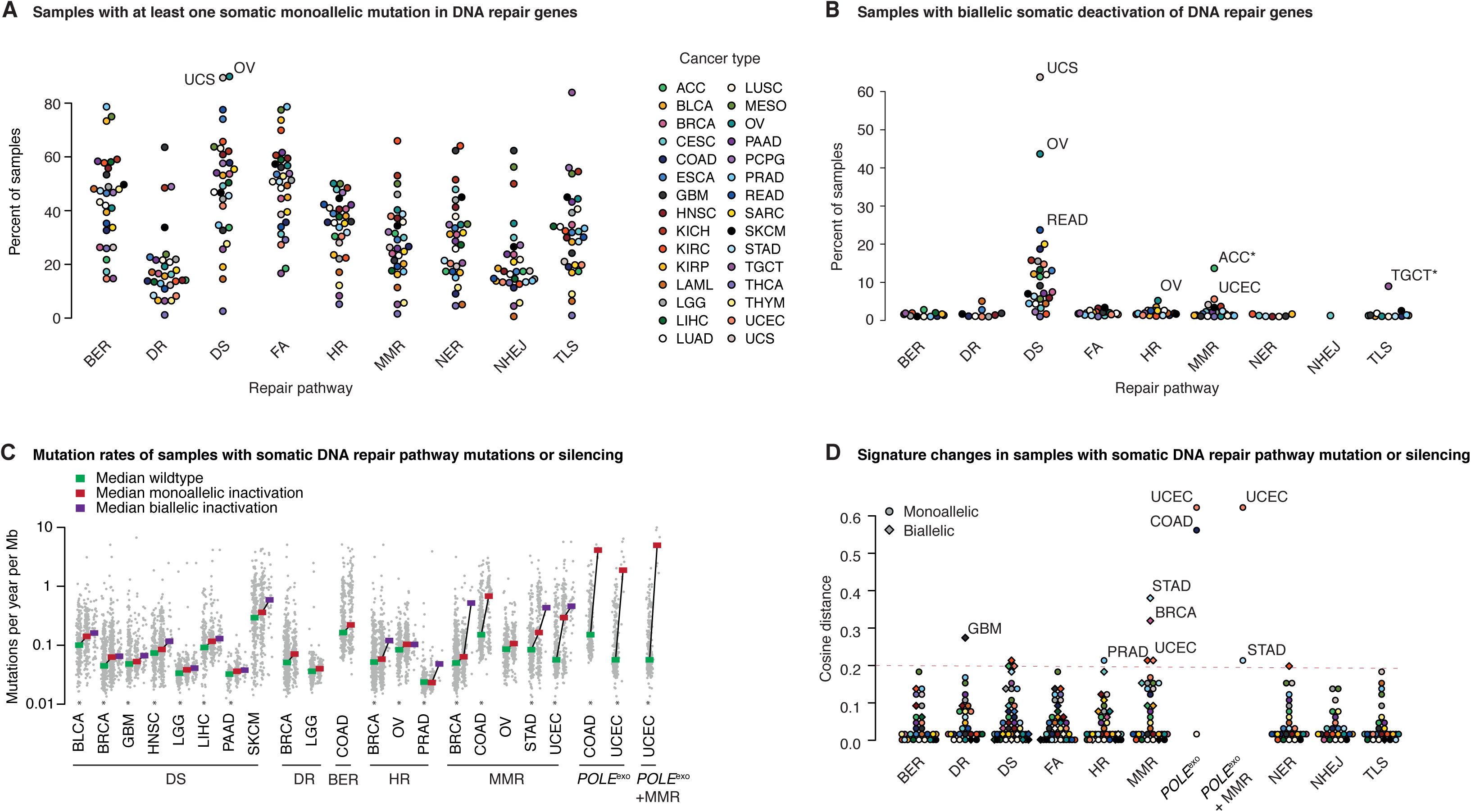
Somatic mutations and silencing of DNA repair pathway genes in human cancers. A. Percentages of samples with mono- or biallelic defects in core components of the DNA repair pathway genes by cancer type for 9,948 cancers from TCGA. B. Percentages of samples with biallelic inactivation of core components of the respective DNA repair pathway by cancer type. C. Age-adjusted mutation rates in samples without (green) or with mono- (red) or biallelic (purple) DNA repair pathway deficiency. Each dot represents the number of mutations per year per Megabase, colored bars denote the medians in each group. Only combinations with Wilcoxon rank sum test FDR under 5% are shown. Stars denote the cancer types where the presence of damaging mutations in the pathway is unlikely to be explained just by the increase in mutational burden (see **Supplementary Methods**). D. Median mutational signature change between samples with DNA repair pathway with respect to median wild-type signature.

**Supplementary Figure 2.**
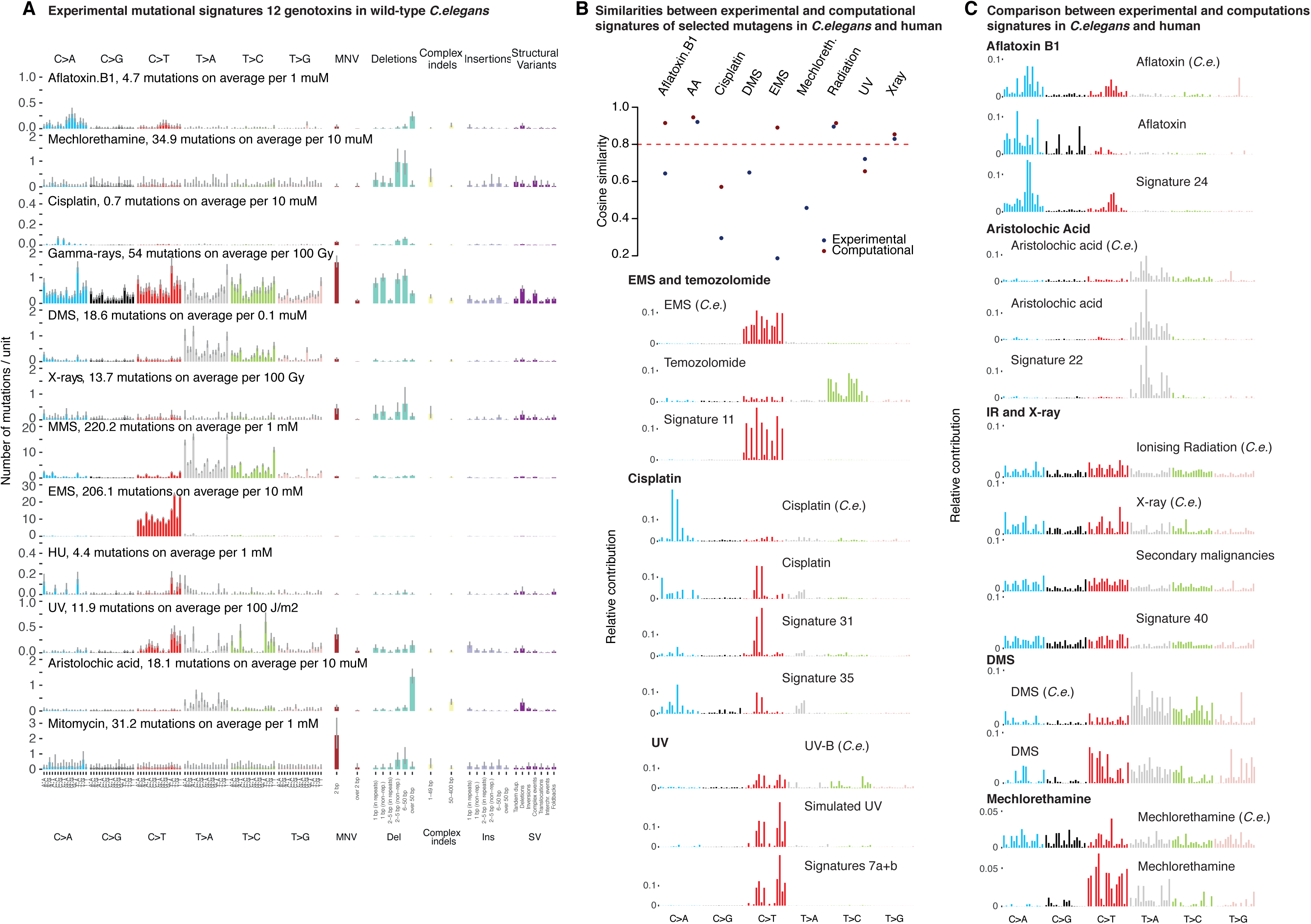
Mutational signatures of genotoxins in wild-type *C. elegans* and their comparison to the experimental mutational signatures of environmental carcinogen exposure^20^, and the catalogue of mutational signatures deduced from tumours^2^. A. Mutational signatures extracted for 12 genotoxins in wild-type *C. elegans.* Each barplot reflects the number of mutations per average dose of indicated mutagen, headers contain the estimate for the total number of mutations per average dose. For details on doses and mutagen specification see **Supplementary Table 1**. For precise numbers of mutations per unit in each class, see **Supplementary Table 2**. B. Cosine similarities between the humanized *C. elegans* signatures and experimental (blue) and computational (red) signatures of same or related mutagens in human cells or cancers. *C. elegans* signatures were adjusted to the human genome trinucleotide frequency. “Experimental” signature for ionizing radiation was estimated as an averaged spectrum of 12 radiotherapy-associated secondary malignancies from ^23^. No counterparts from human data were found for HU, Mitomycin C and MMS. Red line denotes the cut-off of 0.8 above which the spectra are considered similar. C. Visual comparison of experimental signatures for selected mutagens. Signature of aflatoxin was similar to the computationally extracted signature 24 (similarity 0.92) but different from experimental one (0.62). *C. elegans* UV-B exposure showed a C>T mutation spectrum somewhat similar to that in cell line experiments and in cancer, albeit with an additional fraction of T>C mutations. *C. elegans* EMS signature is very different from temozolomide signature in cell lines, but very similar (0.91) to cancer-derived computational signature SBS11 associated with temozolomide. Interestingly, similar phenomena were observed upon exposure with alkylating agents in *Salmonella typhimurium* ^65^. Aristolochic acid signatures were consistent between both experimental systems and cancer. Signature of cisplatin was different from the one identified in cancer or human cell lines. Ionizing radiation and X-rays yielded profiles similar to those in human cancers. Exposures to mechlorethamine and DMS exhibited different mutational spectra in the two model systems.

**Supplementary Figure 3.**
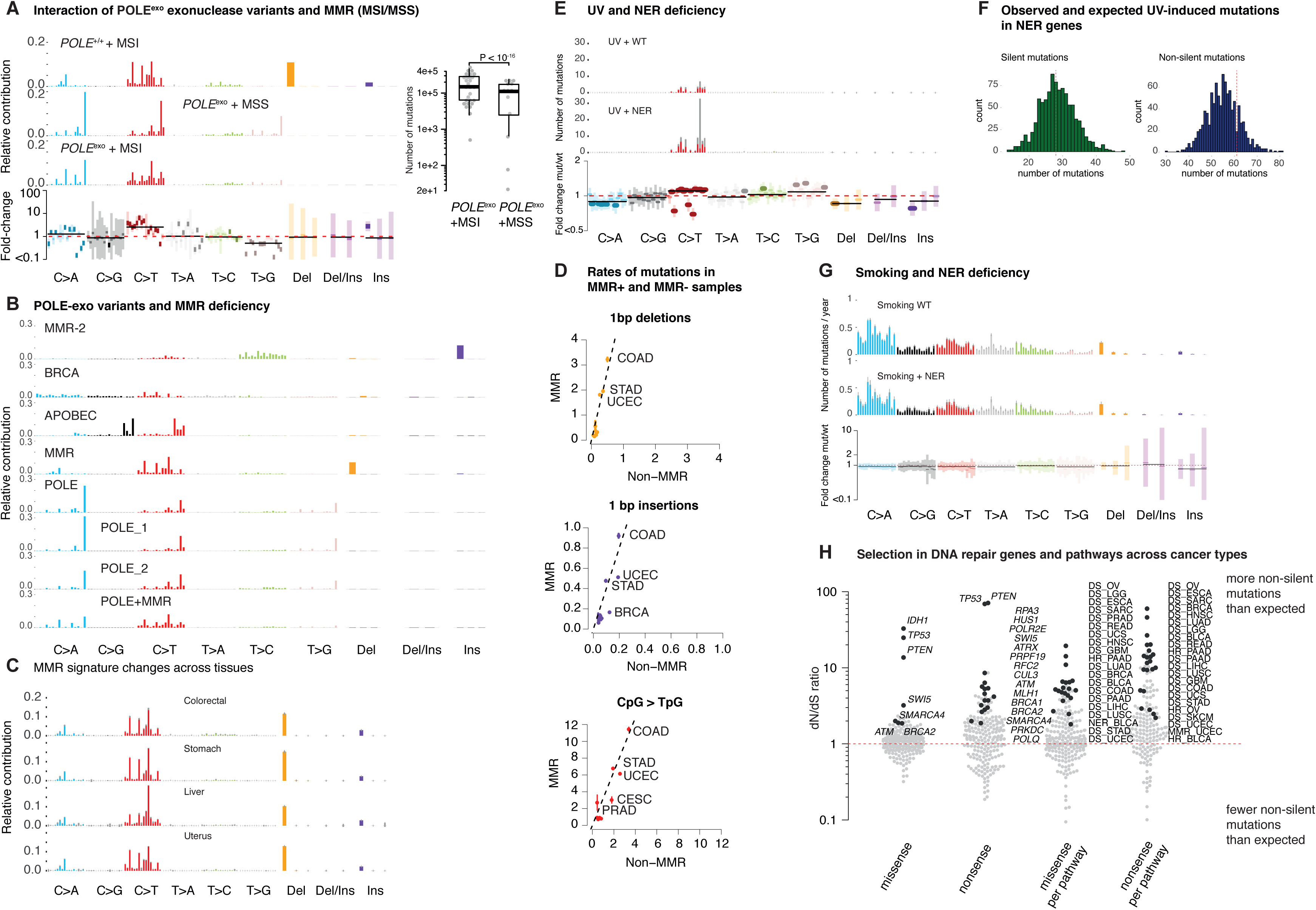
Interaction between damage and repair in human cancer. A. Mutational signatures of MMR deficiency, POLE deficiency, and combined POLE and MMR deficiency. Microsatellite instability is used as a read-out for MMR deficiency. Right: Mutation burden of POLE exonuclease deficiency in MMR proficient (n=40) and deficient (n = 15) uterine cancers. MSI, microsatellite instable; MSS, microsatellite stable. B. 5 mutational signatures extracted from Uterine Cancers, and the POLE exonuclease variant signature with effects of different POLE hotspot mutations (P286R and V411L) and interaction with MMR deficiency (POLE + MMRD). C. Changes of MMR deficiency signatures across different tissues. Barplots represent the appearance of the most widespread MMR signature in different tissues. D. Changes in the rates of specific mutations between MMR proficient and deficient cancers represented by the scatterplot of per-year rates of 1-bp deletions (top), 1-bp insertions (middle) and CpG>TpG mutations (bottom) in tumours with MMR deficiency versus those without it across individual cancer types. Dotted lines represent the linear slope fitted to the rates of MMR deficient versus proficient samples. E. Mutational signature of UV light in melanoma samples with functioning (top panel) and mutated NER (lower panel). The barplot underneath shows the fold-change per mutation type. F. Simulated and observed values for the number of silent and non-silent mutations in NER genes under UV exposure in TCGA SKCM samples with high prevalence of signature 7. G. Mutational signature of tobacco smoking in lung cancer samples with functioning (top panel) and mutated NER (lower panel). The barplot underneath shows the fold-change per mutation type. H. dN/dS values for 248 DNA repair-associated genes across pan-cancer types. dN/dS measures the excess of observed non-silent mutations relative to the expected number based on observed silent mutations. A value of 1 denotes neutrality, while a value greater than 1 signifies a surplus of non-silent changes, indicating a cancer-promoting effect. dN/dS values per gene were calculated separately for missense and nonsense mutations. dN/dS ratios were also calculated for 9 DNA repair pathways (aggregating all genes) across 30 cancer types (right). Black/colored dots reflect results which reached significance compared to the background (FDR 10% within the respective group) with respective genes and DNA repair pathways indicated on the plot. Data for this plot is provided in **Supplementary Tables 4-5**.

**Supplementary Figure 4.**
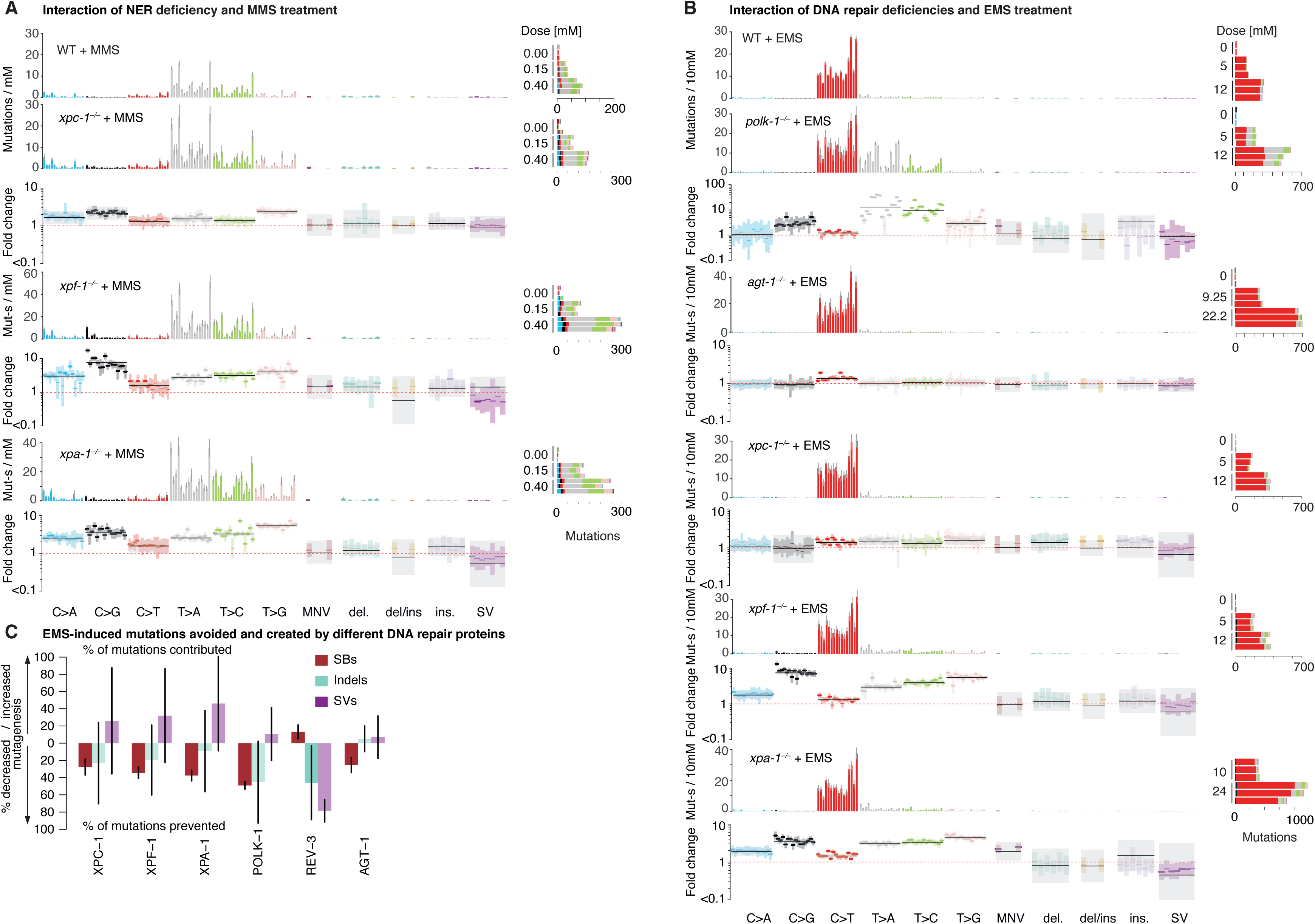
Selected genotoxin-repair interaction cases for alkylating agents in *C. elegans*. A. Mutational signatures and fold-changes per mutation type for MMS exposure in wild-type and NER-deficient backgrounds. Same layout as 3A. B. Mutational signatures and fold-changes per mutation type for EMS exposure in wild-type, *polk-1*^*-/-*^, *agt-1*^*-/-*^ and NER-deficient backgrounds. C. The fractions of mutations contributed or prevented by different DNA repair components compared to the mutations observed upon EMS exposure in wild-type *C. elegans*. Bars filled with more fainted color reflect non-significant changes.

**Supplementary Figure 5.**
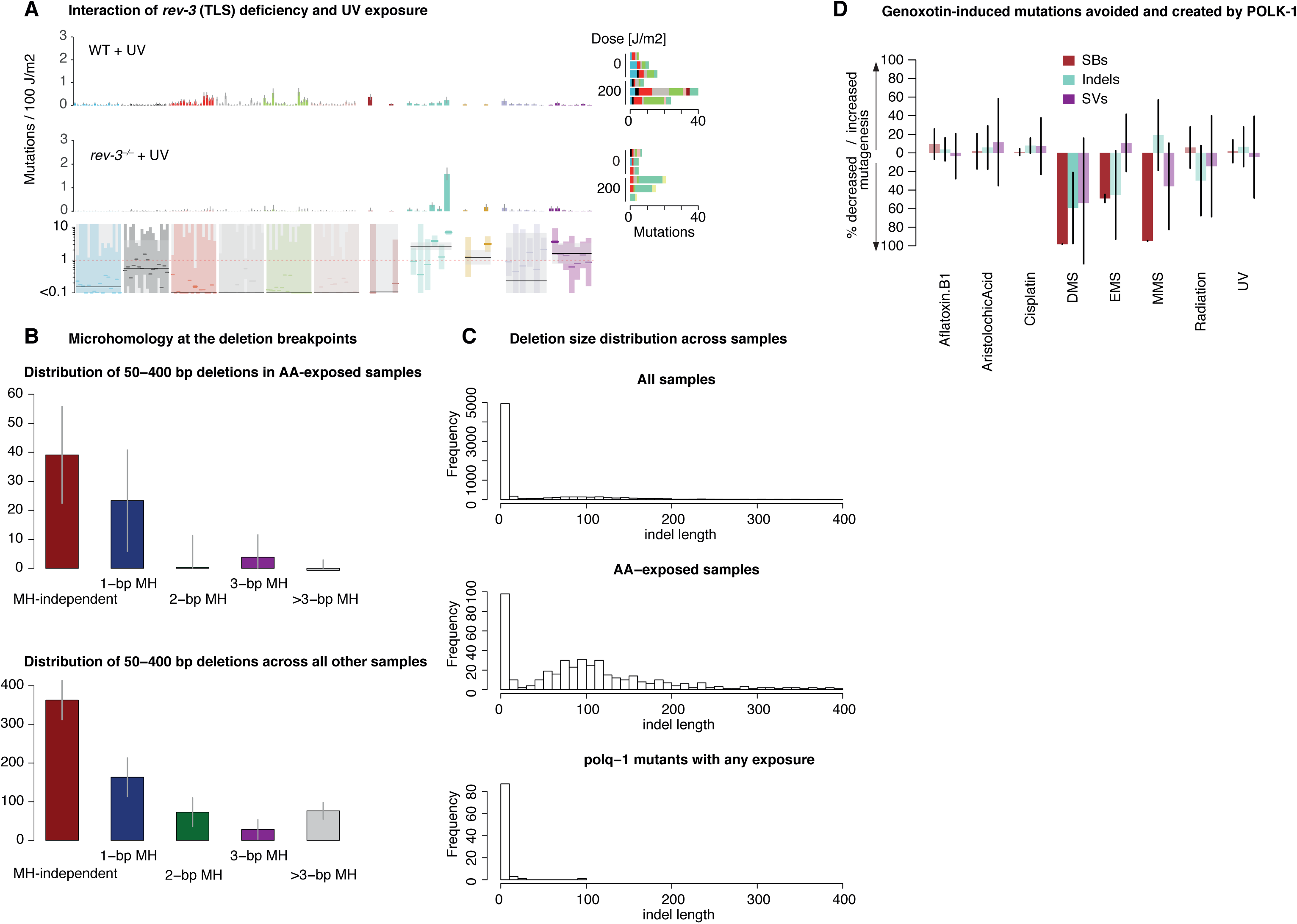
Features of TLS polymerase knock-outs and analysis of deletions in *C. elegans*. A. Mutational signatures and fold-changes per mutation type for UV exposure in wild-type and *rev-3* (TLS) deficient mutants. B. Attribution of 50-400 bp deletions generated in AA-exposed samples (top) and other samples (bottom) based on its microhomology (MH) length. The majority of deletions were likely to occur in a MH-independent manner (red bars). The contribution of deletions with 1-bp MH was substantial in both cases (blue bar), but dominated the spectrum of deletions associated with longer MH in AA-exposed samples (green and purple bars). A high level of short MH indicates the activity of TMEJ contributing to the generation of such deletions across all samples, with an excess of 1-bp MH occuring in AA-exposed samples. C. Distribution of deletion sizes across all samples in the dataset (top), only aristolochic acid exposed samples (middle), and across *polq-1* deficient mutants (bottom). D. The fractions of mutations contributed or prevented by POLK-1 compared to the mutations observed upon exposure to different genotoxins in wild-type *C. elegans*. Bars filled with more fainted color reflect non-significant changes.

**Supplementary Figure 6.**
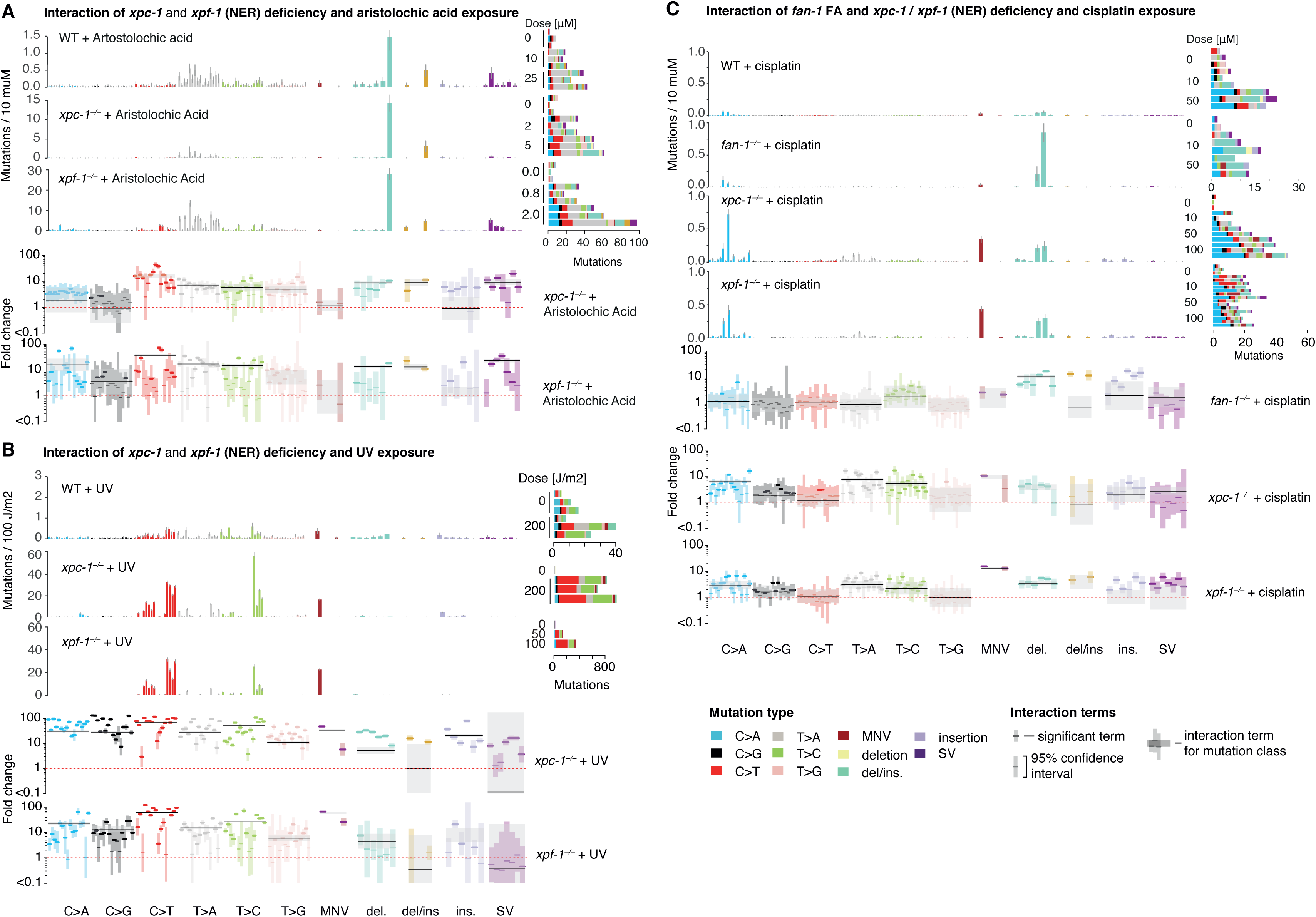
Selected genotoxin-repair interactions involving NER mutants. A. Mutational signatures and fold-changes per mutation type for aristolochic acid exposure under wild-type and NER deficient (*xpc-1* and *xpf-1* mutants) conditions. Note that scale bars are different in the middle and lower panels. B. Mutational signatures and fold-changes per mutation type for UV exposure under wild-type and NER deficient (*xpc-1* and *xpf-1* mutants) conditions. C. Mutational signatures and fold-changes per mutation type for cisplatin exposure in wild-type, under crosslink-repair deficiency (*fan-1* mutants) and NER deficiency (*xpc-1* and *xpf-1* mutants).

## Supplementary Tables

Supplementary Table 1

Description of genetic backgrounds used in the study, sample annotations, mutation counts and design matrix for all *C. elegans* samples.

Supplementary Table 2

Experimental mutational signatures for genetic (including wild-type and DNA repair knockouts) and mutagenic (12 genotoxins used in the study) factors in *C. elegans* along with their 95% credible intervals.

Supplementary Table 3

Genotoxin-repair interaction effects: log fold-changes of mutagen signature per mutation type, and dose-dependent genotype amplification factors along with their 95% credible intervals.

Supplementary Table 4

Map of DNA repair genes with pathways, TCGA cancer names abbreviations, and TCGA samples with mutations in DNA pathways.

Supplementary Table 5

dN/dS values for DNA repair related genes across all cancers types and for DNA repair pathways in different cancer types.

